# A comparative transcriptomic framework across kidney organoid, assembloid, and transplantation models

**DOI:** 10.64898/2026.10.30.762911

**Authors:** Natalija Glibetic, Katie C. Y. Lee, Jason Barnhill, Michael A. Ortega

**Affiliations:** Department of Cell and Molecular Biology, John A. Burns School of Medicine, University of Hawaii, Honolulu, HI

**Author notes:** Address correspondence to: Michael A. Ortega, PhD, Assistant Professor, Department of Cell and Molecular Biology, John A. Burns School of Medicine, 651 Ilalo St. BSB 206, Honolulu, HI 96813.

**Keywords:** kidney, organoids, assembloids, nephron, maturity, scRNA-seq, transcriptomics, benchmark

## Abstract

Human kidney organoids recapitulate key aspects of renal development and have emerged as valuable platforms for disease modeling, nephrotoxicity screening, and therapeutic development. The field now includes a wide range of emerging protocols with distinct cellular compositions and maturation states, which complicates systematic comparison and model selection. Here, we integrated single-cell and single-nucleus transcriptomes from 11 studies spanning kidney organoid and assembloids, both in vitro and following transplantation, benchmarked against fetal and adult kidney references. Transcriptional maturation toward adult identity is compartment specific and does not correlate with protocol complexity. Assembloid protocols show the most adult-like podocyte and distal nephron signatures, transplantation is associated with enhanced tubular oxidative metabolic features, only protocols incorporating ureteric progenitors reliably generate collecting-duct lineages, and no protocol achieves mature adult proximal tubule identity. Annotating against either fetal or adult kidney reference alone leads to misassignment, because early fetal nephron epithelium transcriptionally resembles injury-dedifferentiated adult proximal tubule cells. Assembloid and transplantation models share elevated mechanosensing pathway expression independently of vascularization. Together, these findings provide a framework for kidney organoid model selection.

## INTRODUCTION

Kidney disease affects millions of adults in the United States each year and remains a major contributor to morbidity and mortality^1^. The limited availability of human-relevant experimental systems has been a constraint on mechanistic discovery and therapeutic development in nephrology. Human kidney organoids address this gap by generating multicellular, self-organizing kidney-like structures in controlled *in vitro* environments^2–4^. However, the growing diversity of available protocols has not presented researchers with a systematic cross-protocol approach for selecting the most appropriate model for their experimental needs. Here, we provide a data-driven framework to guide protocol selection across a broad range of applications.

The kidney is a structurally complex organ that develops through three sequential embryonic nephric structures (the pronephros, mesonephros, and metanephros) arising from the intermediate mesoderm. During metanephric kidney development, the ureteric bud (UB) emerges from the mesonephric duct and invades the adjacent metanephric mesenchyme. Although both the UB and metanephric mesenchyme originate from the intermediate mesoderm, they arise from distinct progenitor populations and engage in reciprocal inductive signaling with the UB giving rise to the collecting duct system and the metanephric mesenchyme generating the nephron epithelia. Recapitulating the reciprocal interactions between these two progenitor populations presents a fundamental challenge for in vitro modeling, and as a result most protocols have focused on forming either ureteric or nephron-forming compartments in isolation^5^. The most widely adopted nephron-forming protocols are those of Takasato^6^, Morizane^7^, and Freedman^8^. These protocols generate nephron-like structures containing proximal and distal tubular epithelium and podocytes with varying cellular composition across protocols. However, these protocols also yield heterogeneous stromal populations and limited endothelial cells. Together they provide the current foundation for renal disease modeling. Conversely, ureteric organoid protocols reliably generate branching ureteric structures expressing GATA3, PAX2 and RET in tips and branches. These organoids further differentiate into mature principal cells and developing intercalated cells, yet they lack the nephron-forming components^9–12^. One approach to address these issues has been the development of assembloids^13–17^. These structures combine distinct progenitor populations, such as nephron and ureteric lineages or vascular progenitors, and self-organize to more closely mirror the cellular complexity of the developing kidney. Nevertheless, the transcriptional fidelity and precise developmental positioning of cells within these increasingly complex systems have been difficult to evaluate systematically, underscoring the need for an integrative single-cell analysis to benchmark organoid, assembloid, and transplantation protocols against one another and against *in vivo* fetal and adult kidney references.

One of the first systematic single-cell transcriptomic comparisons of kidney organoids was reported by Wu et al., who profiled organoids generated by the Morizane and Takasato protocols. Both generated a diverse range of kidney cell types, but many were immature and 10-20% were non-renal^18^. Combes et al. subsequently compared organoid and human fetal kidney (hFK) scRNA-seq datasets, while Phipson et al. used bulk and scRNA-seq to examine transcriptional variability within a single protocol across iPSC clones and batches^19,20^. Subramanian et al. profiled over 450,000 cells spanning four iPSC lines across multiple protocols and time points, including organoids transplanted under the mouse kidney capsule^21^. They showed that major cell classes were largely reproducible across iPSC lines, protocols and replicates, and that variability in cell proportions between lines was driven largely by off-target cells. Subramanian et al. also provided the first single-cell characterization of transplanted organoids, showing that transplantation under the kidney capsule reduces these off-target populations. More recently, Yoshimura et al. generated a paired snRNA-seq and snATAC-seq multiome atlas of kidney organoid differentiation from day 7 to 26^22^. Together, these studies characterized the cellular heterogeneity, reproducibility, and differentiation dynamics within or across a limited number of protocols. However, no study has yet integrated nephron organoids, ureteric organoids, assembloids, and transplantation models within a unified transcriptomic analysis.

During normal kidney development, nephron segments mature asynchronously. Podocytes and proximal tubules differentiate early, whereas maturation of the loop of Henle and distal tubule extends into late gestation and the early postnatal period. Maturity is therefore a compartment-specific property. To quantify how far each protocol advances each compartment toward a mature adult state, we integrated single-cell and single-nucleus transcriptomic datasets from 11 studies and benchmarked them against fetal and mature adult kidney references. This comparative framework defines the lineage-specific transcriptional identity and maturity of each model system and provides a practical guide for model selection across a broad range of biological applications.

## RESULTS

### Integration and annotation of fetal, organoid, assembloid, and transplanted organoid kidney tissues

Kidney organoid protocols have considerably diversified, spanning iPSC- and fetal tissue-derived, assembloid, and transplantation-based systems of increasing cellular complexity (**Figure 1**). In this resource, we integrated datasets from 11 separate studies (**Table 1**) to include fetal (weeks 9-18)^23,24^ and adult (KPMP v1.5)^25^ kidney tissue, both iPSC and human fetal tissue-derived nephron-like organoids (Freedman, Morizane, Takasato, and Namestnikov)^18,26,27^, ureteric organoids (Mae)^10^, assembloids (Maggiore, Uchimura, and Huang)^13,14,16^, and organoids transplanted onto the kidney capsule for 30 days (Subramanian)^21^, 60 days (Huang), or subcutaneously for 90 days (Shankar)^28^.

**Figure 1.**
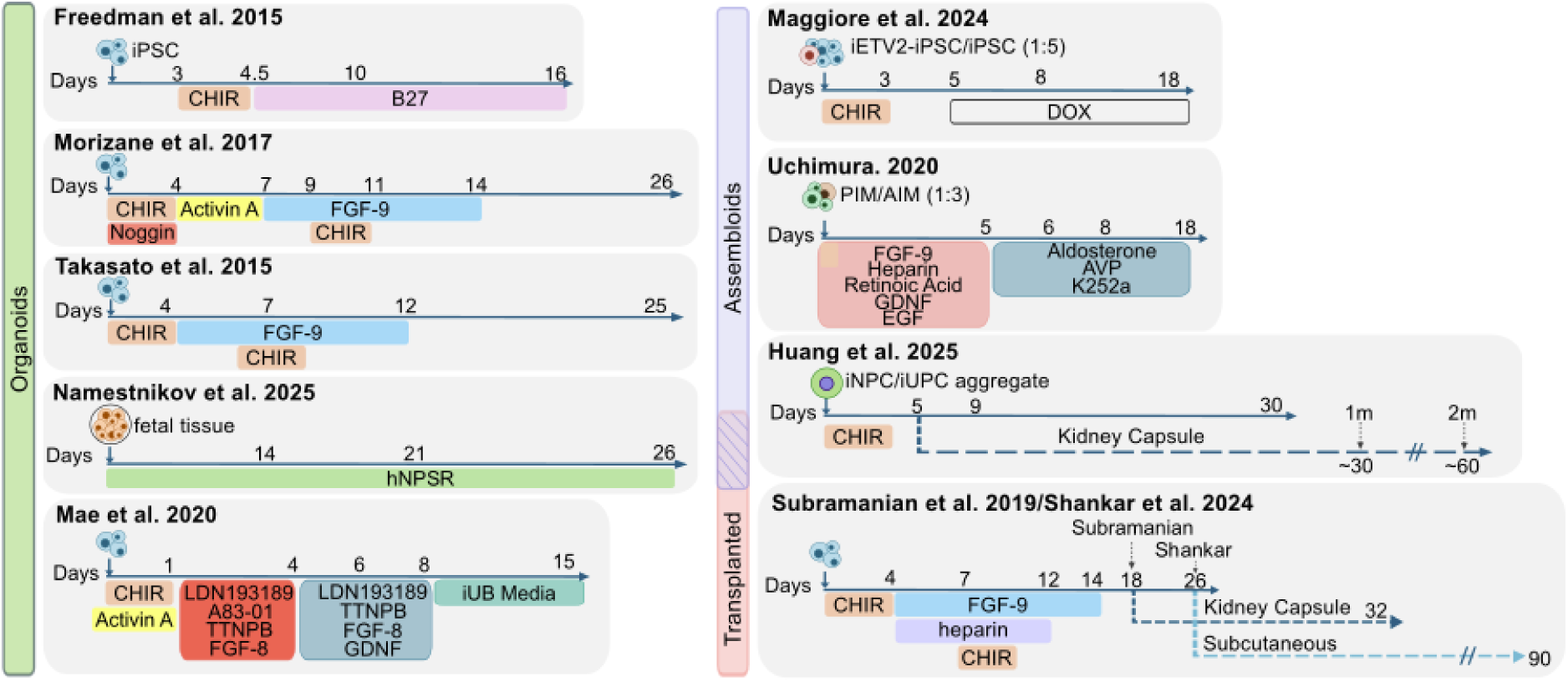
Kidney organoid, assembloid, and transplantation protocols.

**Table 1.**
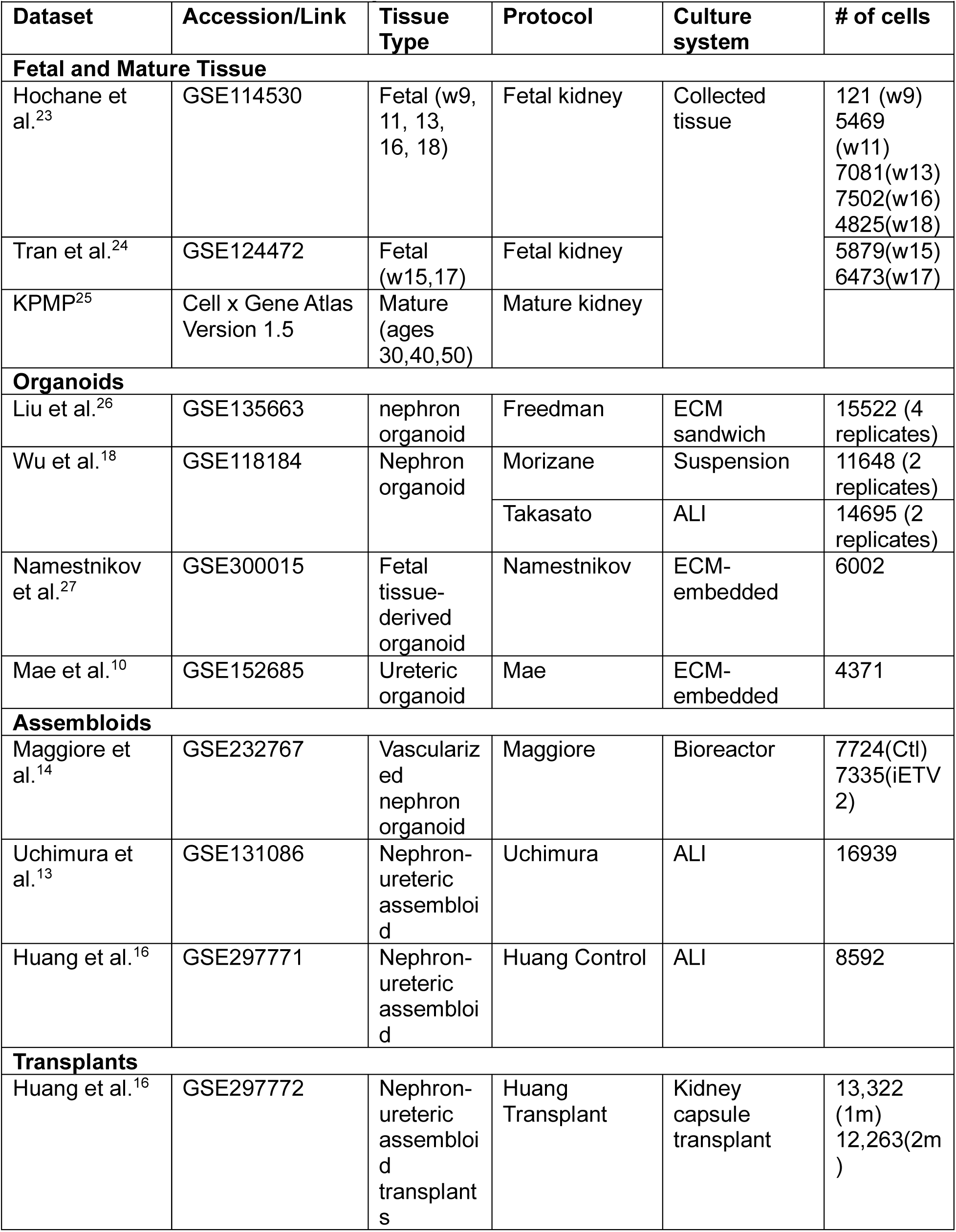
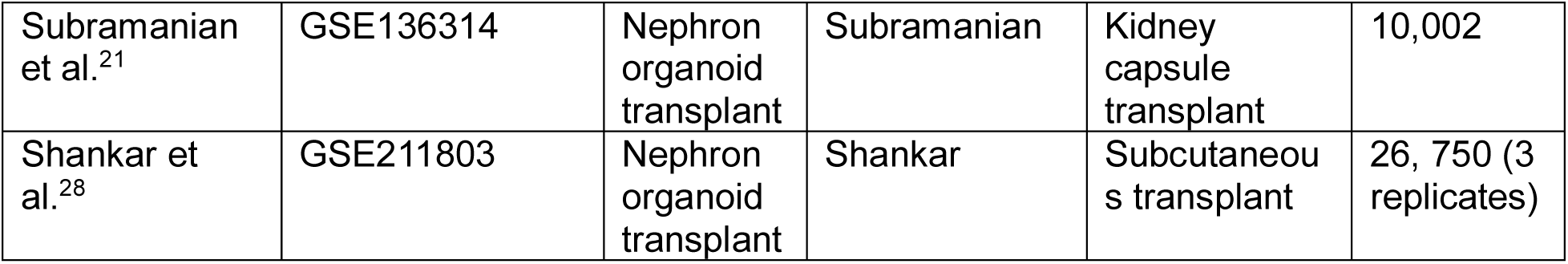
Datasets included in study.

| Dataset | Accession/Link | Tissue Type | Protocol | Culture system | # of cells |
|---|---|---|---|---|---|
| **Fetal and Mature Tissue** |  |  |  |  |  |
| Hochane et al.<sup>23</sup> | GSE114530 | Fetal (w9, 11, 13, 16, 18) | Fetal kidney | Collected tissue | 121 (w9)<br>5469 (w11)<br>7081(w13)<br>7502(w16)<br>4825(w18) |
| Tran et al.<sup>24</sup> | GSE124472 | Fetal (w15,17) | Fetal kidney |  | 5879(w15)<br>6473(w17) |
| KPMP<sup>25</sup> | Cell x Gene Atlas Version 1.5 | Mature (ages 30,40,50) | Mature kidney |  |  |
| **Organoids** |  |  |  |  |  |
| Liu et al.<sup>26</sup> | GSE135663 | nephron organoid | Freedman | ECM sandwich | 15522 (4 replicates) |
| Wu et al.<sup>18</sup> | GSE118184 | Nephron organoid | Morizane | Suspension | 11648 (2 replicates) |
|  |  |  | Takasato | ALI | 14695 (2 replicates) |
| Namestnikov et al.<sup>27</sup> | GSE300015 | Fetal tissue-derived organoid | Namestnikov | ECM-embedded | 6002 |
| Mae et al.<sup>10</sup> | GSE152685 | Ureteric organoid | Mae | ECM-embedded | 4371 |
| **Assembloids** |  |  |  |  |  |
| Maggiore et al.<sup>14</sup> | GSE232767 | Vascularized nephron organoid | Maggiore | Bioreactor | 7724(Ctl)<br>7335(iETV2) |
| Uchimura et al.<sup>13</sup> | GSE131086 | Nephron-ureteric assembloid | Uchimura | ALI | 16939 |
| Huang et al.<sup>16</sup> | GSE297771 | Nephron-ureteric assembloid | Huang Control | ALI | 8592 |
| **Transplants** |  |  |  |  |  |
| Huang et al.<sup>16</sup> | GSE297772 | Nephron-ureteric assembloid transplants | Huang Transplant | Kidney capsule transplant | 13,322 (1m)<br>12,263(2m) |
| Subramanian et al.<sup>21</sup> | GSE136314 | Nephron organoid transplant | Subramanian | Kidney capsule transplant | 10,002 |
| Shankar et al.<sup>28</sup> | GSE211803 | Nephron organoid transplant | Shankar | Subcutaneous transplant | 26, 750 (3 replicates) |

UMAP visualization of the batch-corrected integration shows proper alignment both developmentally and by protocol type (**Figure 2A**). To enable developmental annotation, organoid datasets were integrated alongside fetal kidney references (weeks 9-18), allowing us to anchor transcriptional identities to developmental trajectories (**Figure 2B, C**). Samples were annotated using DevKidCC (DKCC), a cellular identity classification tool trained on fetal kidney tissue data^29^. As expected, cellular composition across individual samples reveals variation in epithelial, stromal, and endothelial proportions, with ureteric epithelium largely restricted to the ureteric progenitor protocols (Mae and Uchimura) and minimal in nephron only organoids and in the Maggiore and Huang assembloids (**Figure 2C**). Off-target cells characterized by SOX2+, STMN2+, and MYOG+ expression (**Figure S1A**)^29,30^ vary considerably across protocols with some organoid (Mae), assembloid (Maggiore) and transplanted samples (Huang) showing substantial contributions from these populations. Consistent with Wilson et al., many of these off-target cells are classified as unassigned or NPC-like, likely reflecting shared expression of nephron progenitor and mesenchymal markers such as SIX2 and, more prominently, CITED1, MEOX1, EYA1 and CRABP2 (**Figure S1B**)^29^. Immune cell markers, including PTPRC+, CCL3+, SRGN+, HLA-DRA+ (**Figure S1C**), were expressed in cells clustering near endothelial populations, alongside fetal and transplanted tissue. Cortical stromal cells were notably enriched in long-term subcutaneous transplants (Shankar) and Matrigel-embedded organoids (Freedman). Endothelial cells are predominantly contributed by fetal samples, with overall low representation across organoid and transplant models. In vascularized assembloids (Maggiore iETV2), this likely reflects a classification artifact rather than true absence as CDH5⁺PECAM1⁺ cells clustering with fetal endothelium were classified as unassigned by DKCC (**Figure 2B**), resulting in underrepresentation of endothelial scores for this protocol.

**Figure 2.**
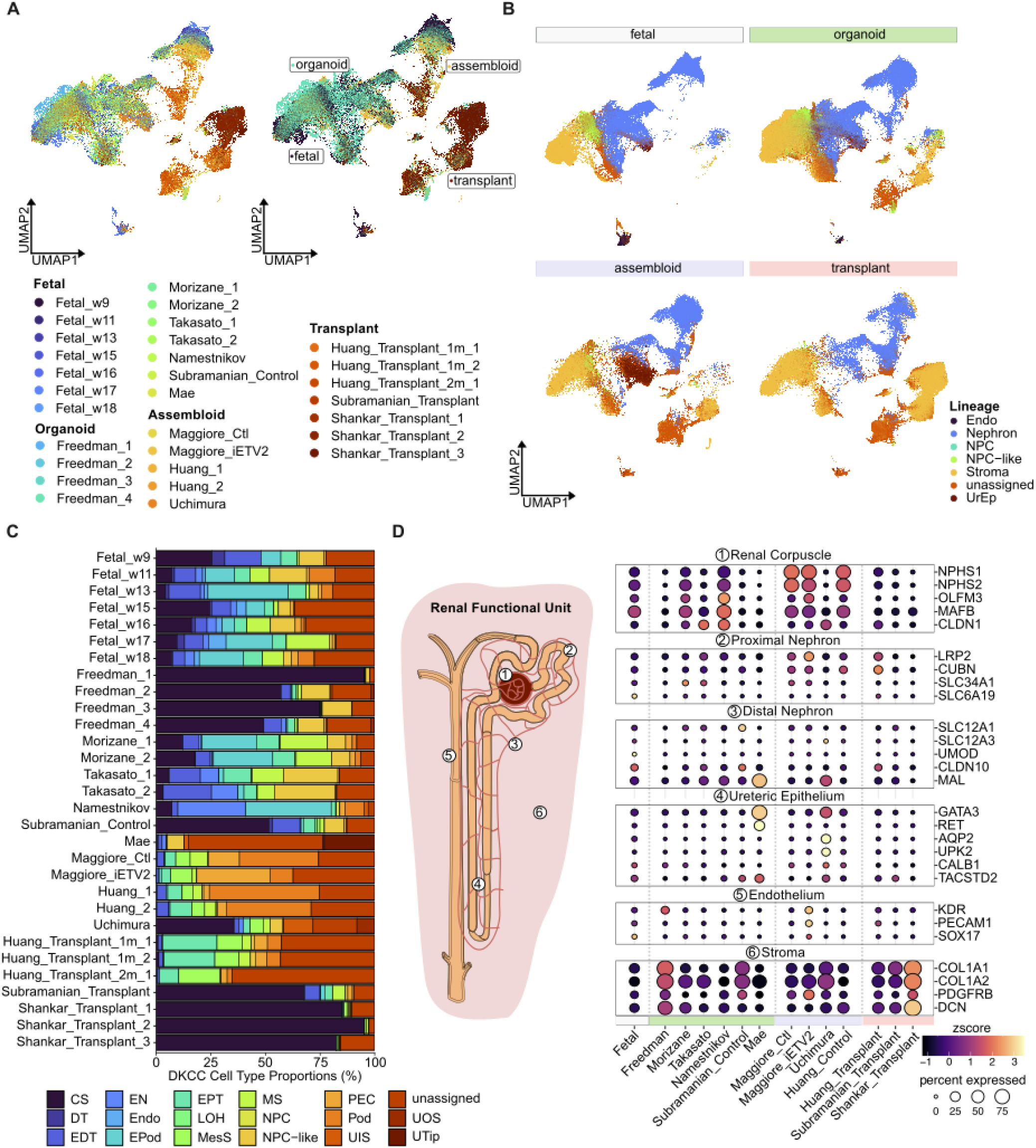
Integrated analysis of fetal, organoid, assembloid, and transplanted organoid kidney tissues. **A)** UMAPs of harmony-integrated data colored by individual sample (left) and by group (right). **B)** UMAP of *DKCC* annotated broad cell lineages. **C)** *DKCC*-annotated cell type proportion by individual sample. **D)** Dotplot comparing the expression of cell type markers for the renal corpuscle, proximal and distal nephron, ureteric epithelium, endothelium, and stroma.

Finally, to validate annotation accuracy, we confirmed segment-specific expression of canonical genes across the renal corpuscle (podocytes, parietal epithelial cells), proximal nephron (proximal tubules), distal nephron (LOH, distal and connecting tubules), ureteric epithelium (inner and outer stalk, ureteric tip), endothelium, and stroma (cortical, medullary, and mesangial cells) (**Figure 2D**). Expression patterns were consistent with expected nephron patterning, supporting DKCC annotations across diverse protocols. Notably, renal corpuscle marker genes for podocytes (NPHS1, NPHS2) were considerably enriched in the Maggiore protocol (both the control and vascularized assembloid), while stromal signatures (COL1A1, COL1A2, PDGFRB, and DCN) were elevated in long-term transplants and ECM-embedded organoids, possibly reflecting substrate-mediated stromal expansion.

### Transcriptional developmental positioning is segment- and protocol-dependent

To assess transcriptional positioning across nephron and ureteric compartments in organoid, assembloid, and transplant systems, we integrated cells from all non-fetal datasets with mature kidney tissue as reference. Initial annotation using a combined KPMP annotation reference for mature, and DKCC reference for non-mature kidney tissue, resolved a broad range of cell types across multiple lineages (**Figure S2**)^25^. To focus the maturity analysis on relevant nephron segments, we subset to nephron lineage populations by removing unassigned, off-target, NPC, NPC-like, immune, and stromal populations. Unsupervised re-clustering resolved 33 transcriptionally distinct cell types spanning the renal corpuscle, proximal nephron, loop of Henle/distal nephron, and ureteric epithelium (**Figure 3A**). Interestingly, integration with the mature kidney reference revealed that the systematic underrepresentation of endothelial cells in DKCC composition scores reflects a misclassification artifact. A transcriptional cluster co-embedding mature kidney endothelial subtypes: glomerular capillary (EC-GC), peritubular capillary (EC-PTC), afferent/efferent arteriole (EC-AEA), ascending vasa recta (EC-AVR), and lymphatic endothelium (EC-LYM), contained 805 assembloid- and transplant-derived cells annotated as unassigned by DKCC. Of these, 419 were transplant-derived, 365 assembloid-derived, and 21 organoid-derived. These unassigned cells expressed canonical endothelial markers PECAM1, CDH5, KDR, and VWF at levels comparable to EC subtypes, confirming endothelial identity (**Figure S3**). Reclassification revealed that Maggiore iETV2 (5.1%), as expected, and Huang transplants (1.7%) harbored the highest endothelial proportions among the protocols, while standard organoids contained substantially lower populations (<1%).

**Figure 3.**
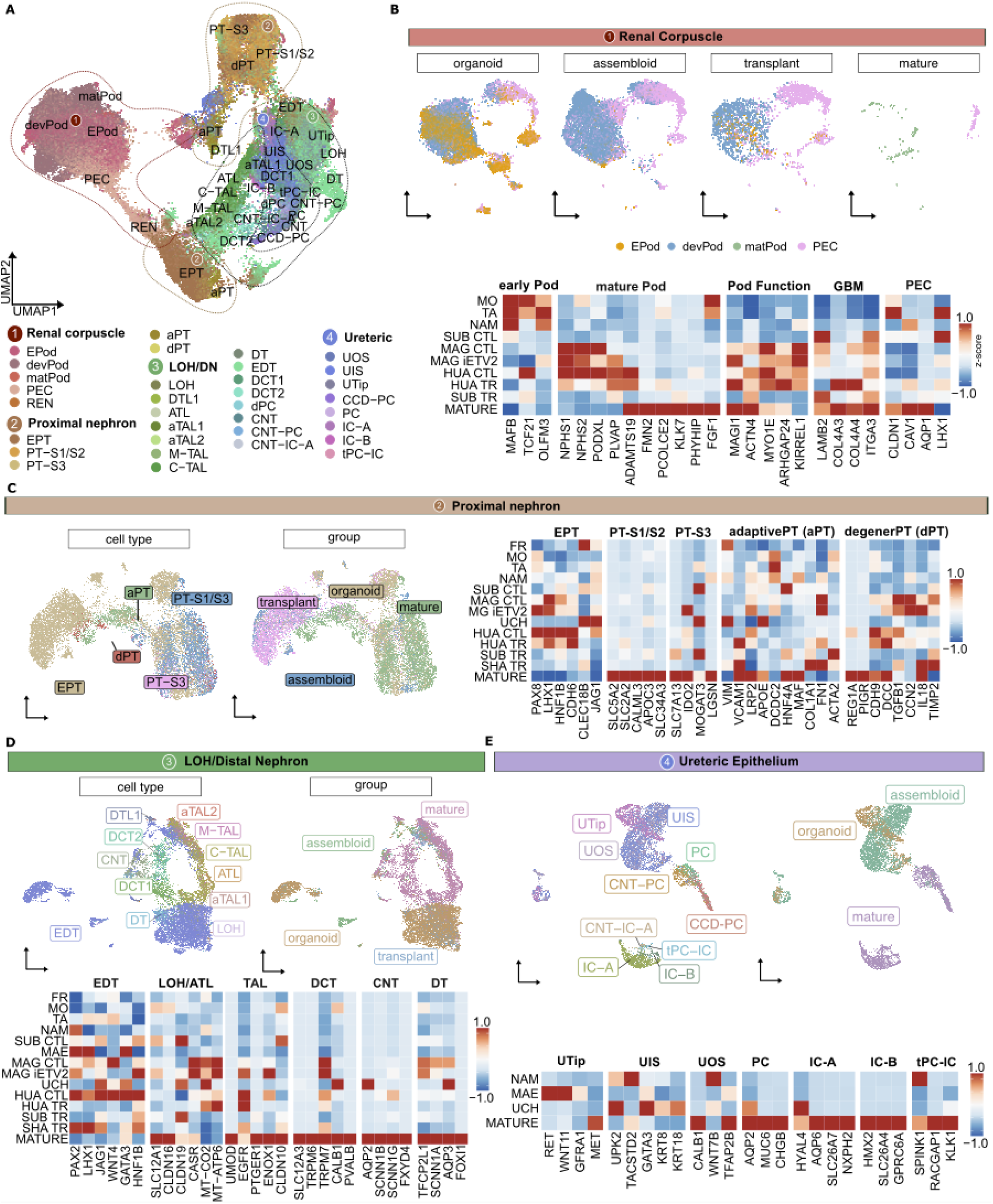
Kidney organoid, assembloid, and transplant models capture distinct nephron populations with varying transcriptional maturity. **A**) UMAP of nephron cells integrated across non-mature and mature tissue colored by cell type. Major nephron compartments are indicated. **B**-**E**) Subclustering of renal corpuscle (**B**), proximal nephron (**C**), LOH/distal nephron (**D**), and ureteric lineage (**E**) populations. For each compartment, UMAPs show cell type distribution across model groups and heatmaps show row-scaled z-scores of curated marker gene expression per protocol.

To examine lineage-specific transcriptional states, we subset and re-integrated cells from four nephron compartments independently: the renal corpuscle, proximal nephron, LOH/distal nephron and ureteric epithelium. Sub-clustering of renal corpuscle cells showed clear differences in podocyte developmental positioning across models (**Figure 3B**). As expected, nephron-like protocols predominantly displayed early podocyte (EPod) transcriptional signatures (MAFB, TCF21, OLFM3), with comparatively lower expression of mature podocyte (Pod) markers, reflecting their fetal-like developmental stage. In contrast, assembloid and transplant models showed a transcriptional shift toward mature podocytes (NPHS1, NPHS2, PODXL) signatures, with near absence of EPod marker expression. However, podocyte markers most enriched in mature tissue in the KPMP atlas (ADAMTS19, FMN2, PCOLCE2, KLK7, PHYHIP, and FGF1) remain predominantly enriched in mature tissue, suggesting that assembloid and transplant protocols achieve broad podocyte maturation but may fall short of the transcriptional identity characteristic of fully differentiated adult podocytes. Beyond individual marker expression, slit diaphragm and foot process genes (MAGI1, ACTN4, KIRREL1, MYO1E and ARHGAP24) were enriched in assembloid podocytes relative to organoids. Glomerular basement membrane (GBM) maturation remained incomplete in assembloids. Assembloid podocytes retained immature LAMB1 but expressed little of mature stage markers COL4A3 and COL4A4. Among all models these collagens were robustly expressed in transplant models. This aligns with Maggiore et al. who reported that vascularized assembloids upregulate COL4A5 but not COL4A3 or COL4A4^14^. The mature α3α4α5(IV) collagen network therefore remains incomplete. Parietal epithelial cell (PEC)-specific markers were most prominent in mature tissue. Gene Ontology (GO) analysis of differentially expressed genes within the renal corpuscle compartment revealed enrichment of glomerular programs, including glomerular filtration and angiotensin maturation, in both assembloid and transplanted assembloid-derived populations (**Figure S4**). Subclustering of proximal tubule populations to early proximal tubule (EPT), S1/S2/S3 segments, mature-specific adaptive proximal tubule (aPT; VCAM1+/DCDC2+), and degenerative proximal tubules (dPT; CST3+/APOE+/ALDOB+), showed a distinct separation between groups with transplants and assembloids co-clustering in one large cluster, and organoid and mature co-clustering separately (**Figure 3C**). EPT gene (PAX8, LHX1, HNF1B, CLEC18B, JAG1) expression was present variably across all organoid and transplant protocols, with highest enrichment in Maggiore and Huang assembloids, but markedly decreased in nephron-like transplants (Subramanian and Shankar). In contrast, S1/S2 (SLC5A2, SLC2A2, CALML3) and S3 (SLC7A13, IDO2, LGSN) were almost exclusively enriched in mature tissue, with canonical segment-specific markers absent across all organoid and transplant models, confirming that neither *in vitro* differentiation nor engraftment is sufficient to drive proximal tubule segment maturation to the level observed in native adult kidney. Interestingly, both adaptive (successful or maladaptive repair) and degenerative proximal tubules overlap significantly with EPTs from non-mature tissue more than with mature S1/S2/S3 proximal tubule segments. Adaptive gene expression (VCAM1, DCDC2, APOE) was detected at moderate levels across all protocols, with VCAM1 expression being notably higher in transplants, while DCDC2 and APOE were higher in organoids. Similarly, dPT markers (CDH9, IL18, DCC, TIMP2) were enriched in both Maggiore and Huang assembloids, as well as Huang transplants, with only TIMP2 higher in Subramanian and Shankar. GO analysis showed that assembloids and transplants showed comparable PT and brush border development, while hFKO and multi-lineage assembloids (Uchimura) display highest metabolic activity across all processes. Consistent with previous observations, mature proximal tubules show preference for fatty acid oxidation and TCA cycle gene programs, while these metabolic pathways were largely absent in most organoid and assembloid protocols, reflecting a more immature metabolic state^31^. However, hFKO and multi-lineage assembloids (Uchimura) show broad upregulation across multiple metabolic programs, suggesting they may engage a more comprehensive metabolic upregulation rather than a lineage-specific or maturation-directed shift (**Figure S4**).

Further subclustering of aPT and EPT populations revealed several overlapping clusters of mature adaptive proximal tubule cells and non-mature early proximal epithelium (**Figure S5A**). Clusters 1 and 2, predominantly composed of mature tissue, showed enrichment for dedifferentiation markers (CD24+, VIM+) (**Figure S5B**)^32–34^. However, these clusters diverged in their functional signatures whereby cluster 1 displayed enrichment for maladaptive markers (VCAM1+, CCL2+, CXCL2+, ACTA2+), while cluster 2 was distinguished by expression of the adaptive marker LRP2+. Cluster 0, the predominantly EPT and non-mature tissue-derived cluster, expressed transitioning/partial EMT markers COL1A1, COL3A1 and FN1 and proliferative markers TOP2A and MKI67 as expected. Clusters 3 and 4 occupied a transcriptionally distinct space, comprising mixed EPT and aPT cell populations with approximately 10% and 25% of cells derived from non-mature tissue, respectively. Cluster 3, the cluster overlapping EPT and mature aPT, was characterized by co-expression of LRP2+, DCDC2+, while cluster 4 exhibited a strongly maladaptive transcriptional signature enriched for markers of cellular transition^35–37^. Notably, the non-mature tissue contributions to these clusters differed by source, where in clusters 0 and 3, non-mature cells were derived predominantly from transplanted Huang assembloids, whereas in cluster 4, they originated from Shankar and Subramanian transplants alongside Morizane organoids.

The distal nephron compartment mirrored proximal tubule findings, with organoid and transplant models predominantly harboring transcriptionally immature distal cells (**Figure 3D**). Regardless of annotation source, canonical markers of mature distal nephron segments including UMOD for TAL, SLC12A3 and TRPM6 for DCT, and AQP2 and SCNN1B for CNT, were largely absent across organoid and transplant protocols, with expression confined almost exclusively to mature tissue. The TAL transporter SLC12A1 was, however, detected at reduced levels across several organoid and transplant protocols (23% of organoid and 18% of transplant distal cells, up to 38% in Subramanian control) rather than absent. A notable exception was the Uchimura assembloid model, which showed elevated expression of AQP2, AQP3, CALB1, CLDN19, JAG1, and GATA3, markers spanning collecting duct, CNT, and ureteric lineages. This is consistent with Uchimura’s protocol incorporating both nephron and ureteric-like components. Notably, despite comparable cell numbers, Huang transplants showed markedly reduced expression of both EDT and mature distal nephron markers relative to their non-transplanted assembloid counterpart, yet displayed elevated expression of mitochondrial oxidative phosphorylation genes MT-CO2 and MT-ATP6 suggesting transplantation may drive these cells into a transcriptionally intermediate but metabolically altered state. Importantly, this was consistent across all transplant models, none of which showed appreciable expression of mature distal segment markers UMOD, SLC12A3, TRPM6, or SCNN1B, potentially suggesting engraftment alone is insufficient to drive distal nephron segmental maturation.

While nephron compartment maturation varied systematically across protocols, the ureteric compartment showed the most restricted distribution, with ureteric identity largely confined to a subset of models (**Figure 3E**). Ureteric tip (UTip) was most highly expressed in Mae, consistent with its ureteric bud differentiation protocol. Ureteric inner stalk (UIS) markers were most prominent in Uchimura and hFKOs, while ureteric outer stalk (UOS) cells were present mainly in Uchimura, at low levels in Mae, and not detectable in hFKOs. Principal cell (PC) markers, including AQP2, were detected in Uchimura but were largely absent across all other organoid protocols, and expression of the transitional PC-IC marker SPINK1 was most prominent in hFKOs. These findings reflect the differences in protocol design where Mae and Uchimura explicitly incorporate ureteric progenitors, while hFKOs, as a fetal tissue-derived model, captures a distinct subset of ureteric lineages, likely reflecting the developmental stage of the donor tissue.

Together, these analyses show that protocols vary in the lineages they generate as well as in the maturation of those lineages. The Maggiore control and iETV2 assembloids were largely indistinguishable across the nephron segment maturation analysis, differing mainly in higher expression of basement membrane markers (COL4A5, LAMA5, LAMC1) and CCN2 in the iETV2 model.

### Adult kidney atlas transfer resolves lineage-specific classification gaps across developmental and transplant models

During compositional analysis, we noted that Huang assembloid, despite incorporating both nephron and ureteric progenitors, showed complete absence of ureteric epithelium by DKCC. This prompted us to perform a complementary label transfer using the KPMP adult kidney atlas as a reference, which enabled orthogonal classification of cell types beyond the developmental window captured by DKCC. This approach revealed systematic discordances between the two methods that were both protocol- and lineage-dependent (**Figure 4**) and provided biological justification for reclassifying populations DKCC did not resolve.

**Figure 4.**
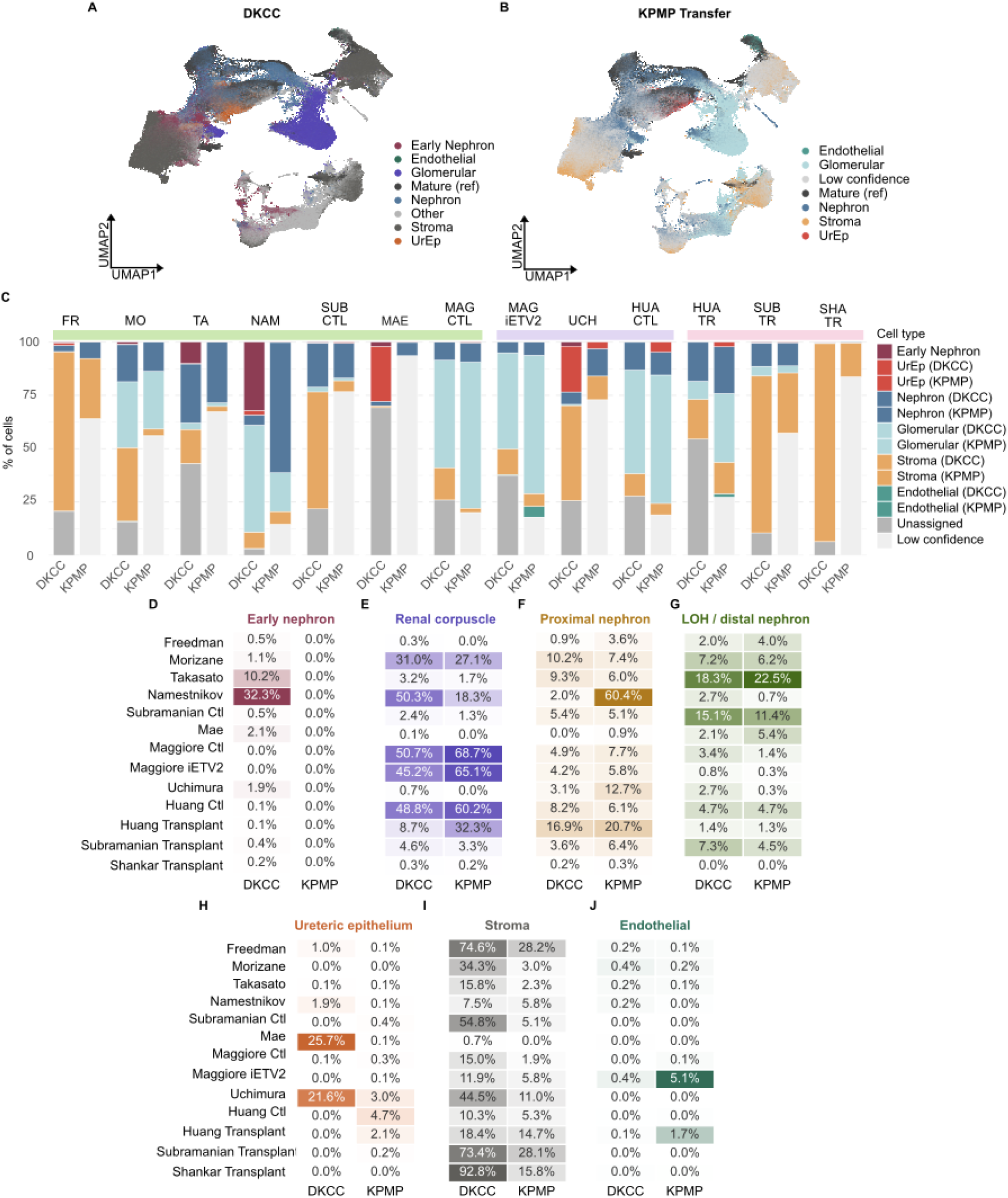
Integrated UMAP by DKCC cell type annotation (**A**) and by KPMP adult kidney atlas label transfer (**B**). Grey indicates low-confidence predictions (score <0.5). **C**) Cell type composition per protocol by DKCC (left) and KPMP transfer (right). (**D-J**) Percentage of cells per protocol assigned to early nephron (**D**), renal corpuscle (**E**), proximal nephron (**F**), LOH/distal nephron (**G**), ureteric epithelium (**H**), stroma (**I**), and endothelium (**J**) by DKCC and KPMP transfer. Color scale is independent per lineage to maximize within-lineage contrast.

For the early nephron, DKCC identified EN cells predominantly in Namestnikov hFKOs (30.8%) and Takasato organoids (10.2%), with low representation across other protocols analyzed (**Figure 4C, D**). As expected, KPMP transfer resulted in zero EN signal across all protocols, confirming this progenitor state has no direct equivalent in the adult kidney atlas and is exclusively captured by the developmental reference.

For the ureteric lineage, DKCC identified early ureteric populations (UTip, UIS, UOS) predominantly in Mae (25.7%) and Uchimura (21.6%), consistent with their ureteric bud strategies. KPMP transfer recovered collecting duct epithelium (principal cells and connecting tubule) almost exclusively in the Huang assembloid and transplant protocols (4.7% and 2.1%, respectively), and Uchimura (3.0%) with negligible signal across the remaining protocols. Together, these two methods capture complementary stages of the ureteric lineage trajectory, with DKCC anchoring the progenitor end and the KPMP transfer resolving the differentiated end.

For the proximal nephron, the most striking finding was observed in Namestnikov hFKOs. DKCC assigned only 3.6% of cells to proximal nephron progenitor states (NPC, NPC-like, and EPT), while the KPMP transfer identified 60.4% as transcriptionally proximal (**Figure 4**). Closer inspection revealed that this signal is driven almost entirely by EN (early nephron, 30.8%) and EPod (early podocyte, 39.2%) cells mapping specifically to dedifferentiated adaptive proximal tubule (aPT; CD24+, VIM+, APOE+) state in KPMP rather than to mature S1/S2/S3 proximal tubule segments (**Figure S6A, B**). Module scoring confirmed both EN (1.68) and EPod (0.99) cells strongly express the dedifferentiated aPT gene signature, with negligible adaptive (LRP2) or maladaptive (VCAM1, DCDC2) signatures (**Figure S6C**). This places fetal nephron epithelium and injury-dedifferentiated adult proximal tubule in shared transcriptional space despite their fundamentally different biological origins. Notably, 26.7% of EPod cells correctly mapped to mature podocytes, suggesting a subset achieved sufficient maturity. Conversely, Takasato, Morizane and Freedman organoids showed the inverse pattern, with high DKCC proximal progenitor signal (35.6%, 18.6%, and 10.6%) but low KPMP transfer agreement (6.0%, 7.4%, and 3.6%), respectively. This may reflect retention of an early nephron progenitor state that has not yet committed to a mature tubular identity. For the distal nephron, agreement between annotations was generally closer, with all protocols showing broadly consistent signals. This suggests that distal nephron identity is more consistently recognized across both the fetal developmental reference and the adult reference. A notable exception was Uchimura, where DKCC assigned only 2.7% to distal nephron while KPMP transfer assigned only 0.3%, consistent with Uchimura’s enrichment to ureteric rather than distal nephron lineages, which may share surface marker expression but are transcriptionally distinct at the level of the adult atlas.

For the glomerular compartment, the KPMP transfer consistently recovered higher proportions than DKCC in Maggiore and Huang protocols. Huang transplants (8.7% DKCC vs 32.3% KPMP transfer) and Maggiore control assembloid (50.7% vs 68.7% and iETV2 45.2% vs 64.9%). In these protocols, a larger fraction of podocyte compartment cells were assigned to adult glomerular identity (POD, PEC, and PL) by KPMP than to the developmental renal corpuscle by DKCC, consistent with our subclustering analysis above. For stroma, DKCC substantially overassigned stromal identity in long-term transplant models relative to the KPMP transfer, with Subramanian and Shankar transplants reaching 73.4% and 92.8% by DKCC compared to only 28.1% and 15.8% by KPMP, respectively.

Finally, for the endothelial lineage, KPMP transfer provided direct biological justification for reclassifying a population of 805 assembloid- and transplant -derived cells annotated as unassigned by DKCC. These cell co-embed with mature kidney endothelial subtypes: glomerular capillary (EC-GC), peritubular capillary (EC-PTC), afferent/efferent arteriole (EC-AEA), ascending vasa recta (EC-AVR), and lymphatic endothelium (EC-LYM) (**Figure S3A, B**). These cells expressed canonical endothelial markers PECAM1, CDH5, and KDR at levels comparable to KPMP endothelial subtypes, supporting their reclassification as endothelium (**Figure S3C**). This reclassification revealed that Maggiore iETV2 (5.1%) and Huang transplants (1.7%) harbored the highest endothelial proportions among non-mature protocols.

Taken together, DKCC and KPMP-based label transfer provide complementary and largely concordant annotations, with DKCC more comprehensive in resolving the developmental progenitor states of each lineage and KPMP assignment of the more mature adult-like states. In both the proximal and ureteric lineages, however, the mature identities KPMP assigned, dedifferentiated proximal tubule in Namestnikov and collecting duct in Huang and Uchimura were carried by progenitor-retaining epithelium. The cells assigned to proximal tubules retained the nephron progenitor factors PAX8 and PAX2 but lacked mature proximal segment genes, and those assigned to collecting duct were GATA3+ but lacked FOXI1 and mature principal cell genes. Both therefore reflect transcriptional positioning toward adult identity rather than robust maturation.

### Mechanosensing pathway activity varies systematically across protocols

Physical forces, including fluid dynamics, tissue tension, compression, and pressure, have emerged as key regulators of tissue maturation, yet their contribution to protocol-specific differences in kidney organoid nephron identity remains poorly understood^38–40^. To investigate whether protocol-specific differences in nephron maturation are accompanied by differences in mechanosensing, we examined the transcriptional expression of established mechanotransduction pathways across protocols (**Figure 5**). Analysis of the Hippo pathway, a key mechanosensing regulator of tissue maturation, organ size and homeostasis, revealed distinct protocol-specific patterns^41–43^. Huang and Maggiore assembloids showed elevated YAP1 (YAP) expression along with Hippo kinases MST2, LATS1, and LATS2, while canonical YAP targets CYR61 and AMOTL2 remained low (**Figure 5, left panel**). Together, this indicates that elevated YAP and HIPPO pathway expression in these assembloids was not accompanied by induction of these canonical YAP targets. Maggiore and Huang also showed elevated primary cilia flow sensing genes PKD2, IFT88, and INVS (**Figure 5, middle left panel**), and Maggiore assembloids additionally showed elevated integrin-focal adhesion genes ITGB1, PTK2, and VCL (**Figure 5, middle right panel)**. Meanwhile, the Maggiore control and iETV2 were almost indistinguishable across these genes, with the vascularized iETV2 model modestly higher.

**Figure 5.**
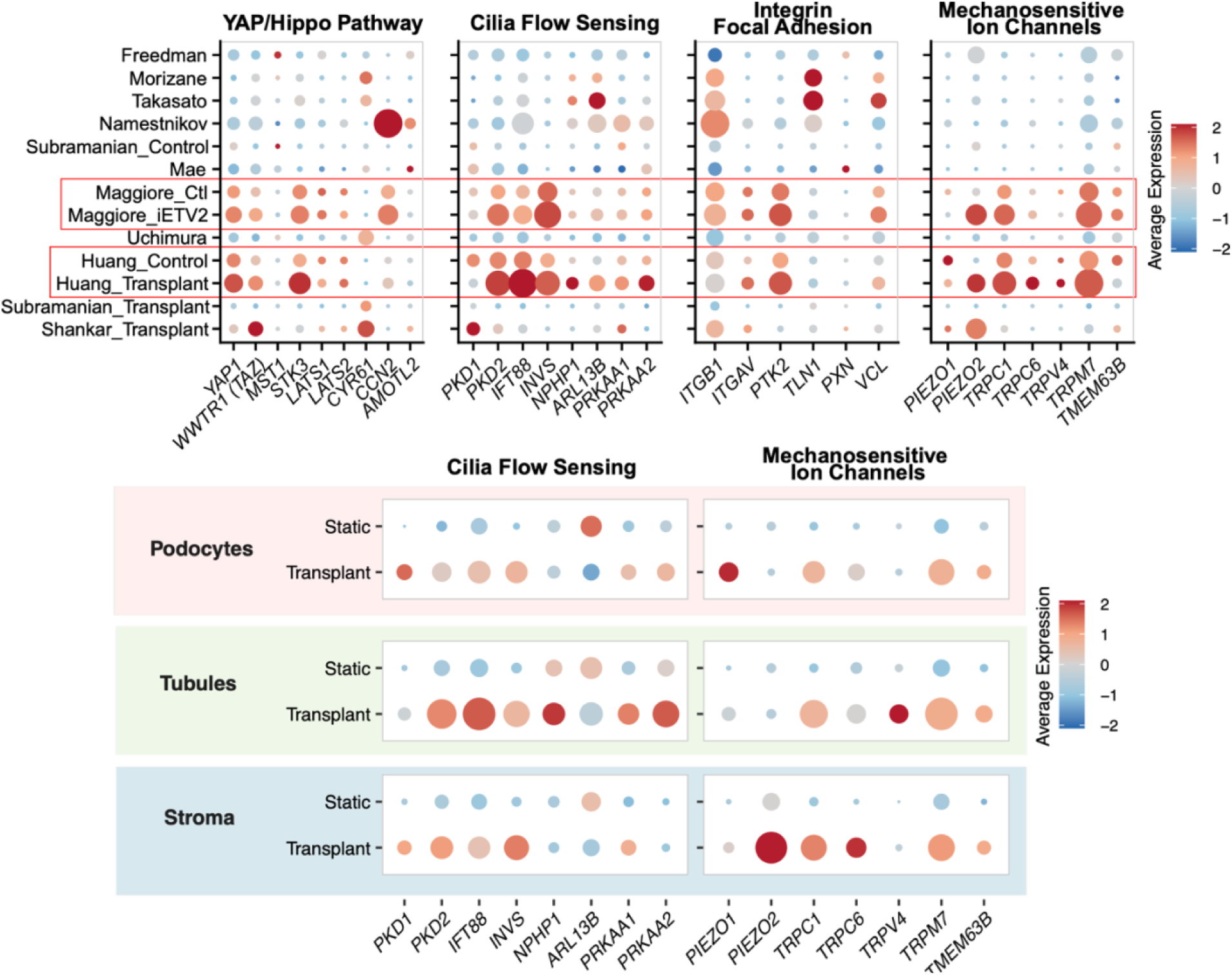
Mechanosensing pathway signatures vary across protocols. Dot plots showing scaled expression of Hippo pathway (**left**), cilia flow-sensing (**middle left**), and integrin/focal adhesion (**middle right**), and mechanosensitive ion channel (**right**) genes across organoid, assembloid and transplant models. Dot size indicates proportion of cells expressing each gene, while color indicates scaled mean expression. Expression of the cilia flow and mechanosensitive ion channel gene sets within matched podocyte, tubular, and stromal compartments of the Takasato organoid (static) and transplanted Huang assembloid.

Analysis of mechanosensitive ion channel genes further showed elevated TRPC1 and TRPM7 across Maggiore and Huang relative to other protocols, and elevated PIEZO2 in the vascularized and transplanted models (Maggiore iETV2, Huang transplant, and Shankar transplant), while PIEZO1, TRPV4, and TMEM63B remained low across protocols (**Figure 5, right panel**). Across the remaining protocols these mechanotransduction transcripts were uniformly low, with the lowest representation in the nephron-only organoids and in the ureteric progenitor protocols Mae and Uchimura. The long-term subcutaneous Shankar transplant was a partial exception, showing elevated canonical YAP target genes CYR61 and AMOTL2 together with PIEZO1 and PIEZO2 rather than the upstream YAP1 and Hippo kinase pattern seen in the assembloids. Together, these mechanosensitive signatures indicate that mechanotransduction transcript expression differs systematically across protocols. Here, we find these signatures to be elevated in the Maggiore, Huang, and the Shankar transplant models, while found to be lowest in the nephron-only and ureteric progenitor organoids.

Because the Huang transplant contains a larger podocyte fraction than the Takasato organoid, 8.7% compared to 3.2% of cells, elevated mechanosensing expression measured across all cells could reflect a difference in cell type abundance rather than cell specific signatures. To determine which cell populations carried these mechanosensing signatures, and to contrast a static organoid with a vascularized transplant, we compared the mechanosensitive ion channel and primary cilia gene sets within matched compartments of the Takasato organoid (static) and the transplanted Huang assembloid. Within podocytes, tubular epithelium, and stromal cells, both gene sets were higher in the transplant than in the static organoid model by roughly 5 to 6-fold for the ion channel signatures and 2 to 3-fold for the primary cilia signatures. The signatures largely localized to the compartments in which these mechanosensors are normally expressed, with the primary cilia flow-sensing genes PKD2, IFT88, and NPHP1 together with the mechanosensitive channel TRPV4 highest in tubular epithelium, PIEZO1 highest in podocytes, and PIEZO2 highest in stroma. These compartment specific signatures suggest higher expression in the transplant reflects intrinsic cellular expression rather than a difference in podocyte abundance.

## DISCUSSION

The diversity of human kidney organoids has posed a challenge for researchers seeking models appropriate for their biological question. While individual protocols have been validated for specific applications, and previous single-cell studies compared a small number of nephron organoid protocols and iPSC lines, this study provides a systematic comparison spanning organoid, assembloid, and transplantation models, establishing a data-driven resource for protocol selection.

The integrated dataset presented here reveals that kidney organoid protocols differ primarily in which segments achieve advanced transcriptional developmental positioning. These distinctions have direct and non-obvious implications for model selection that cannot be inferred from structural complexity or protocol alone. Annotation against fetal kidney references confirmed conserved nephron and stromal identities across all protocols, with ureteric epithelium restricted to the ureteric progenitor protocols Mae and Uchimura (**Figure 2**). Off-target neural and muscle populations were present and varied across most protocols. While targeted interventions and transplantation offer partial relief, none eliminate them entirely^18,21,44^. Recent single-cell transcriptomic trajectory analysis of kidney organoid differentiation revealed their origins in incomplete mesodermal patterning during primitive streak induction suggesting that full elimination may require more precise anterior-posterior specification at the earliest stages of differentiation rather than downstream protocol adjustments^30^. Ureteric epithelium appears only in dedicated ureteric-lineage protocols Mae and Uchimura, whereas the Huang assembloid, despite incorporating ureteric progenitors, yielded no developmental ureteric epithelium. This indicates that incorporating ureteric progenitors is not by itself sufficient and that the dual-lineage architecture of the kidney cannot be replicated by nephron differentiation alone.

Developmental classification alone, however, proved insufficient to capture the full cellular landscape of more transcriptionally advanced protocols. DKCC, trained on fetal kidney references, correctly resolved ureteric progenitors (UTip, UIS, UOS) in Mae and Uchimura, and early nephron epithelium (EN) most prominently in Namestnikov hFKOs (30.8%) and Takasato (10.2%), reflecting the enrichment in progenitor populations in these protocols. However, DKCC failed to detect ureteric epithelium in Huang assembloids despite their explicit dual lineage design, and systematically left mature populations unresolved in transplant and assembloid conditions. Complementary label transfer using KPMP adult kidney atlas recovered collecting duct epithelium in Huang assembloids and transplants, and reclassified 805 endothelial cells missed by DKCC in Maggiore iETV2 and Huang transplants. These recovered collecting duct cells were GATA3+ but lacked FOXI1 and mature principal cell genes, indicating ureteric lineage positioning without principal or intercalated cell maturation (**Figure 4**). Moreover, it revealed that apparent proximal tubule signal in Namestnikov hFKOs reflects mismapping of early nephron and early podocyte cells to a dedifferentiated adaptive proximal tubule state in KPMP, a transcriptional signature shared between fetal progenitor epithelium and injured-dedifferentiated adult proximal tubule (**Figure 4, Figure S6**). Because the adult adaptive proximal tubule is itself a dedifferentiated PAX2+ and CD133+ progenitor-like state, and the organoid cells expressed this signature without maladaptive markers VCAM1 and DCDC2 (**Figure S6**), the overlap likely reflects developmental immaturity rather than any unintended injury during induction^25^. Thus, in both the proximal and ureteric lineages, the adult reference assigned mature identities to cells still carrying progenitor programs. Pairing the adult reference with a developmental one can therefore reveal confounding assignments.

Segment-resolved benchmarking against adult kidney tissue showed that protocol advantages in transcriptional positioning are compartment specific rather than global (**Figure 3**). Rather than more complex assembloid and transplant models outperforming simpler organoids, we observe each protocol advanced different segments and none advanced them all. The renal corpuscle showed the clearest maturation. Assembloid protocols showed the most advanced podocyte and distal nephron signatures, yet no protocol achieved full adult-like proximal tubule identity. By contrast, proximal tubule metabolic programming was elevated in a subset of protocols independent of this segment identity. Specifically, most protocols generated cells co-expressing early proximal markers rather than committed S1 to S3 identity, consistent with the field-wide recognition that proximal tubule maturation represents a central bottleneck, even in protocols designed to address it^5,45–47^. Notably, while canonical segment-specific proximal tubule identity remains absent across all systems, hFKOs and the multi-lineage Uchimura assembloid display broad upregulation across oxidative phosphorylation, fatty acid oxidation, TCA cycle, and ion transport programs (**Figure 3C**). However, this upregulation in hFKOs and the Uchimura assembloid seemingly occurred without segment-specific transporters, indicating that metabolic capacity and segment maturation are separable in these systems. This distinction has direct implications for how functional maturity of kidney organoids is assessed. Across all protocols, ureteric epithelium retained early transcriptional programs without progression toward mature principal or intercalated cell identity, thus collecting duct maturation likewise remains unresolved. That the Huang transplant and Maggiore assembloids share elevated mitochondrial respiratory chain signatures (MT-CO2, MT-ATP6) suggests a metabolically altered state in distal tubular epithelium common to *in vivo* engraftment and bioreactor culture. Interestingly, these two conditions also share improved oxygenation relative to static culture, a plausible basis for the oxidative shift that warrants further investigation.

Mechanosensing enrichment was convergent across the Maggiore and Huang assembloids and the Shankar long-term subcutaneous transplant (**Figure 5**). In the assembloids this enrichment paralleled their more advanced podocyte and tubular epithelial signatures. The Hippo signaling pathway, primary cilia-AMPK signaling, and integrin-focal adhesion mechanotransduction were each elevated in these protocols relative to other organoids. The most consistent of these signals was the induction of mechanosensitive ion channels, with TRPC1 and TRPM7 elevated across Maggiore and Huang models. Interestingly, the similar transcriptional signatures between Maggiore control and vascularized assembloids suggest a shared mechanical variable rather than vascularization. This interpretation is supported by a growing body of evidence that mechanosensing is broadly a regulator of organoid maturation^48^. Homan et al. (2019) showed that flow-induced shear stress expanded endothelial progenitor populations and generated perfusable vascular networks, accompanied by enhanced podocyte polarity and adult tubular gene expression compared to static controls^49^. Hiratsuka et al. (2022) showed that fluid flow engages RAC1 and FOS mechanosensing in the distal nephron of kidney organoids-on-a-chip i a polycystic kidney disease model where this stimulus drove cyst formation^50^. Finally, Tekguc et al. demonstrated that stirred bioreactors promote glomerular vascularization through mechanosensory integrin α2β1 signaling, with coordinated upregulation of ECM organization and actin cytoskeletal remodeling, which is in agreement with the integrin-focal adhesion enrichment identified in Maggiore and Huang assembloids^51^. Whether these mechanosensory differences contribute to maturation or simply mark the physically distinct culture environments of these protocols remains to be tested directly.

Translating these findings into practice, this resource (**Figure 6**) supports that protocol selection should be application-driven, with assembloid systems favored for glomerular and tubular biology, nephron organoids retained for accessible PT nephrotoxicity screening, and transplantation models reserved for applications requiring *in vivo* maturation at the cost of complexity. Beyond marker-based classification and single-reference annotation strategies, combining DKCC for developmental progenitor resolution with KPMP-based label transfer for mature cell type identification offers a more complete characterization of protocols and should be considered when annotating any kidney organoid system where developmental stage is uncertain or heterogeneous. Only protocols incorporating ureteric progenitors generated collecting duct lineage, and even these remained short of mature principal or intercalated cell identity. Here, several limitations warrant acknowledgment. This study relies on previously generated datasets varying in protocol, sample composition, and sequencing depth. While batch correction was applied, residual technical variability may remain. Several recent approaches to enhance organoid maturity or structural organization were not included, as the underlying data are not publicly available or were generated under conditions precluding direct comparison. A key gap is the absence of publicly available human kidney references spanning late fetal, neonatal and pediatric stages. Developmental maturity was therefore anchored to fetal and mature adult tissue, leaving intermediate stages unresolved. Transcriptomic similarity to mature kidney, while principled, does not directly capture transport kinetics or filtration function. Even the assembloid podocytes that approached mature marker expression retained immature LAMB1 with little COL4A3 or COL4A4 and an unassembled α3α4α5(IV) network. Nevertheless, this resource represents a comprehensive segment-resolved cross-protocol comparison, and provides a practical evidence-based foundation for kidney organoid model selection.

**Figure 6.**
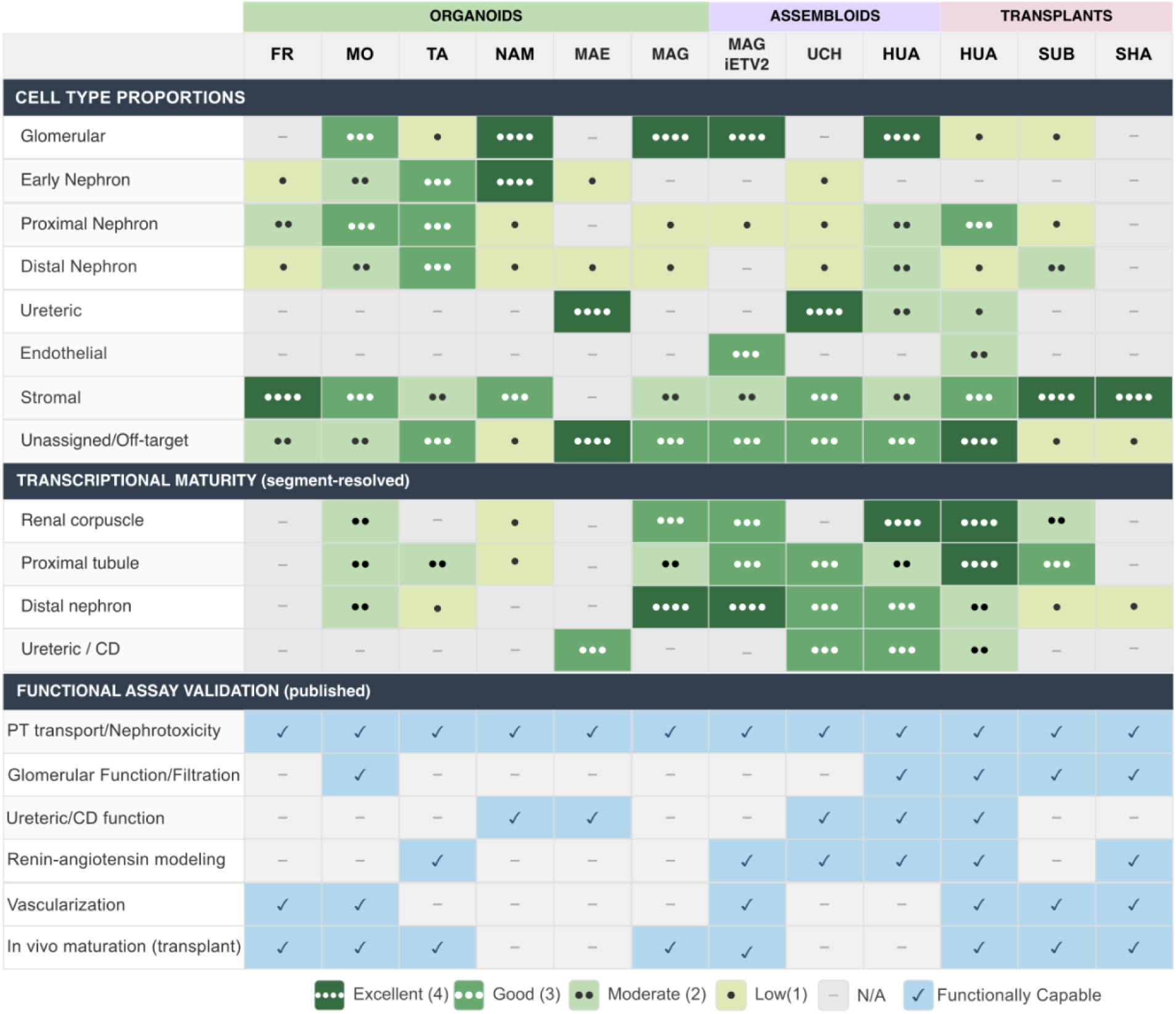
Protocol selection framework for kidney organoid applications. Comparative scoring matrix summarizing cell type composition, transcriptional maturity, and published functional assay validation across organoid, assembloid, and transplantation protocols. Cell type proportions and transcriptional maturity scores are derived from the integrated single-cell dataset; functional assay checkmarks reflect published demonstrations in the literature.

## Methods

### Experimental Model and Subject Details

No new human subjects or experimental animals were recruited for this study.

### Datasets

Publicly available single-cell and single-nucleus RNA-seq datasets were collected from 13 sources (see **Table 1**). Reference datasets comprised human fetal kidney (weeks 9–18) and the Kidney Precision Medicine Project (KPMP v1.5) healthy adult kidney atlas. Query datasets were grouped as organoids (Freedman, Morizane, Takasato, fetal kidney tissue-derived Namestnikov, and Mae ureteric organoids), assembloids (Maggiore control and iETV2-vascularized; Uchimura and Huang nephron-ureteric assembloids), and transplants (Huang kidney capsule, Subramanian kidney capsule, and Shankar subcutaneous). All datasets were obtained as processed count matrices from the original deposits.

### Preprocessing and quality control

Before merging, each dataset was preprocessed to harmonize gene nomenclature and remove dataset-specific artifacts. This included removing reference-genome prefixes from gene symbols (e.g., hg19-, GRCh38-) and removing empty droplets with DropletUtils. For transplanted organoid datasets aligned to combined human–mouse references, mouse-derived features were removed. For the fetal comparisons, all samples except mature were merged into a single object and quality control thresholds were set per sample type. Quality control thresholds were set per sample type: fetal tissue samples were filtered to retain cells with > 300 detected genes and < 10% mitochondrial reads, whereas organoid, assembloid, and transplant samples were filtered at < 20% mitochondrial reads. For all samples cells were retained if 200 < nFeature_RNA < 6,000, and nCount_RNA < 40,000. One sample (Huang_Transplant_2m_2) was excluded entirely due to markedly reduced library quality relative to its paired replicate (Huang_Transplant_2m_1), with a median of 374 genes and 481 UMIs per cell compared to 1,049 and 1,572 respectively. Quality control metrics were summarized before and after filtering to ensure consistency across samples. Following quality control, the fetal comparison object retained 213,520 cells across 30 individual samples, comprising 37,350 fetal, 80,967 organoid, 32,866 assembloid and 62,337 transplant cells. For the mature comparisons, all samples except fetal were merged and processed in the same way. Following final quality control, the mature comparison object retained 194,414 cells were retained across 39 individual samples, comprising of 18,244 mature, 80,967 organoid, 32,866 assembloid and 62,337 transplant cells. The merged dataset was log-normalized, highly variable genes (3000 for fetal-anchored, and 2000 for mature-anchored objects) were identified, and the scaled data reduced by principal component analysis (PCA; 50 PCs). Before integration, UMAP embeddings were generated using the first 30 PCs to visualize dataset-specific structure and batch effects.

### Harmony integration and clustering

For fetal-anchored integration, all organoid, assembloid, transplant, and fetal kidney datasets were merged. Data were log-normalized, the top 3,000 highly variable genes were identified, and the scaled data were reduced by PCA (50 PCs). Sample-level batch effects were corrected using Harmony^52^ with sample as the grouping variable and default parameters. The first 20 Harmony dimensions were used for nearest-neighbor graph construction (k = 20), Louvain clustering (resolution 0.5), and UMAP visualization. For maturity-focused integration, fetal datasets were excluded, and the same integration pipeline was applied to organoid, assembloid, transplant, and mature kidney datasets, using the top 2,000 highly variable genes (HVG).

### Cell type annotation

Cell type identity was assigned using *DevKidCC* (DKCC), a reference-based classifier trained on human fetal kidney single-cell data, which assigns lineage identity labels to each cell^29^. For mature kidney cells, author-provided cell type annotations from the KPMP v1.5 atlas were used directly. To identify cell types beyond the developmental window captured by DKCC, labels were transferred from the mature kidney cells of the KPMP v1.5 dataset (author annotations^25^) to all organoid, assembloid and transplant cells. The reference was reprocessed (3,000 HVGs, scaling, PCA with 30 PCs). Transfer anchors were identified with FindTransferAnchors on log-normalized data using the reference PCA, and labels were transferred using TransferData (dimensions 1-30). Cells with a maximum prediction score ≥ 0.5 received the transferred label. All others were classified as low confidence. Predicted labels were grouped into broad lineage categories (proximal nephron, distal nephron, renal corpuscle, ureteric epithelium, stroma, and endothelium). To quantify lineage representation, the percentage of cells per protocol assigned to each lineage was computed independently for DKCC and KPMP transfer, using lineage-to-cell type mappings across both methods. DKCC lineage groupings used were early nephron (EN, NPC), renal corpuscle (Pod, EPod, PEC), proximal nephron (EPT), LOH/distal nephron (LOH, DT, EDT), ureteric epithelium (UOS, UIS, UTip), stroma (CS, MesS, MS), and endothelium (Endo). KPMP transfer lineage groups used were proximal nephron (PT-S1/S2, PT-S3, aPT, dPT, cycEPI), distal nephron (DTL1, ATL, C-TAL, M-TAL, aTAL1, aTAL2, DCT1, DCT2), renal corpuscle (POD, PEC, PL), ureteric epithelium (PC, CCD-PC, dPC, tPC-IC, IC-A, IC-B, CNT-IC-A, CNT, CNT-PC), stroma (FIB, dFIB, MyoF, VSMC, VSMC/P, MC, dVSMC, REN, aFIB), and endothelium (EC-GC, EC-PTC, EC-AVR, EC-AEA, EC-LYM). A combined annotation was generated merging DKCC labels and KPMP annotations with manual curation of ambiguous populations. Endothelial cells in certain protocols were identified by co-clustering with mature kidney endothelial subtypes and confirmed by expression of canonical markers (PECAM1, CDH5, KDR, VWF). These 805 cells were reclassified as endothelium.

### Segment subclustering

To perform segment-resolved analyses, cells were subset by broad lineage into four compartments: renal corpuscle (early podocytes, mature podocytes, parietal epithelial cells, and juxtaglomerular renin-expressing cells), proximal nephron (early proximal tubule, proximal segments S1/S2, S3, adaptive proximal tubule, and degenerative proximal tubule), distal nephron (loop of Henle, descending and ascending thin limb, medullary and cortical thick ascending limb, distal convoluted tubule, connecting tubule and early distal tubule), ureteric epithelium (ureteric tip, inner stalk, outer stalk, principal cells, intercalated cells, and transitional principal-intercalated cells). Renin-expressing cells were assigned to the renal corpuscle by juxtaglomerular anatomical association for the segment-resolved analysis, whereas the lineage-composition analysis grouped them with stroma following the KPMP developmental-lineage convention. In addition, the stromal segment included cortical, medullary, mesenchymal stroma, fibroblasts, myofibroblasts, and vascular smooth muscle cells, and the endothelial segment included cells sub-clustering together with early endothelium, glomerular capillary (EC-GC), peritubular capillary (EC-PTC), afferent/efferent arteriole (EC-AEA), ascending vasa recta (EC-AVR), and lymphatic endothelium (EC-LYM) and unassigned cells. Differentially expressed genes were identified using FindAllMarkers in Seurat with a Wilcoxon rank-sum test, retaining genes expressed in at least 25% of cells within a cluster with a log2 fold change threshold of 0.5. Gene Ontology enrichment analysis was performed on cluster marker gene lists using standard overrepresentation testing.

### Pseudobulk-based transcriptional maturity scoring

Segment-resolved transcriptional maturity was assessed by computing pseudobulk average expression of curated mature kidney marker gene sets within each nephron compartment, restricted to cells of the corresponding lineage. Marker genes were selected based on literature-confirmed cell type markers and enrichment in the KPMP v1.5 atlas. For each compartment, z-scored pseudobulk expression was averaged across marker gene groups to generate a per-protocol maturity score. Protocols were ranked relative to one another within each compartment. Mature kidney was included as a reference but excluded from inter-protocol ranking.

### Mechanosensitive pathway analysis

To assess mechanotransduction pathway activity across protocols, curated gene sets were defined for three pathways: YAP/Hippo signaling (YAP1, WWTR1, LATS1, LATS2, CYR61, CTGF/CCN2, AMOTL2, NF2), primary cilia-AMPK signaling (PKD1, PKD2, IFT88, INVS, NPHP1, ARL13B, PRKAA1, PRKAA2, SIRT1), and integrin-focal adhesion mechanotransduction (ITGA2, ITGB1, ITGAV, VCL, TLN1, PTK2, PXN, ROCK1, ROCK2, RAC1, ACTA2, COL4A1, LAMA5, LAMB2, LAMC1), and PIEZO/TRP mechanosensitive channels (PIEZO1, PIEZO2, TRPC1, TRPC6, TRPV4, TRPM7, TMEM63B). For each gene set, pseudobulk average expression was computed per protocol across all cells, z-scored across protocols, and visualized via heatmap.

### Visualization

All visualizations were generated in R using ggplot2, Seurat, SeuratExtend, and patchwork. Heatmaps of pseudobulk expression used a blue-white-red color scale with z-scoring applied across protocols within each gene group. UMAP embeddings, DotPlots, and proportion bar charts were generated using SeuratExtend. Figures were created in Inkscape v1.4.2

## RESOURCE AVAILABILITY

### Lead Contact

All requests for additional information and resources should be directed to Michael A. Ortega

### Materials availability

No new reagents and materials were used in this study.

### Data and code availability

This paper analyzes existing, publicly available data. Accession numbers for all datasets are listed in Table 1, and the software packages used for single-cell processing are listed in the key resources table. This paper does not report original code. Any additional information required to reanalyze the data reported in this paper is available from the lead contact upon request.

## Key Resources Table

| **Reagent or resource** | **Source** | **Identifier** |
|---|---|---|
| **Software and algorithms** |  |  |
| R v[4.4.2] | R Core Team | https://www.r-project.org |
| Seurat v[5.4.0] | Hao et al., 2021<sup>53</sup><br>Hao et al., 2024<sup>54</sup> | https://satijalab.org/seurat/ |
| Harmony v[1.2.4] | Korsunsky et al., 2019<sup>55</sup> | https://github.com/immunogenomics/harmony |
| DropletUtils v[1.26.0] | Lun et al., 2019<sup>56</sup> | https://bioconductor.org/packages/DropletUtils |
| DevKidCC v[0.4.0] | Wilson et al., 2022<sup>29</sup> | https://github.com/KidneyRegeneration/DevKidCC |
| SeuratExtend v[1.2.7] | Hua et al., 2025<sup>57</sup> | https://github.com/huayc09/SeuratExtend |
| AUCell v[1.28.0] | Aibar et al. 2017<sup>58</sup> | https://bioconductor.org/packages/AUCell |
| GSVA v[2.0.7] | Hänzelmann et al., 2013<sup>59</sup> | https://bioconductor.org/packages/release/data/annotation/html/org.Hs.eg.db.html |
| Msigdbr v[26.1.0] | Dolgalev, 2026<sup>60</sup><br>Liberzon et al., 2015<sup>61</sup> | https://cran.r-project.org/package=msigdbr |
| edgeR v[4.4.2] | Chen et al., 2025<sup>62</sup> | https://bioconductor.org/packages/edgeR |
| org.Hs.eg.db v[3.20.0] | Carlson, 2024<sup>63</sup> | https://bioconductor.org/packages/org.Hs.eg.db |
| ggplot2 v[4.0.2] | Wickham, 2016 | https://ggplot2.tidyverse.org |

## ACKNOWLEDGMENTS

This work was funded by NIH NIDDK R21DK133754 and NIGMS P20GM113134 to M.O. and P20GM103466 to J.B.

## AUTHOR CONTRIBUTIONS

N.G. and M.A.O. conceived the study. N.G. performed data collection and analysis. K, C.Y.L, and J.B. contributed to data analysis and study coordination. M.A.O. supervised the project. N.G. drafted the manuscript. N.G., K.C.Y. L., J.B. and M.A.O. revised the manuscript.

