## Supplemental Figures for "A comparative transcriptomic framework across kidney organoid, assembloid, and transplantation models"

### SUPPLEMENTAL INFORMATION

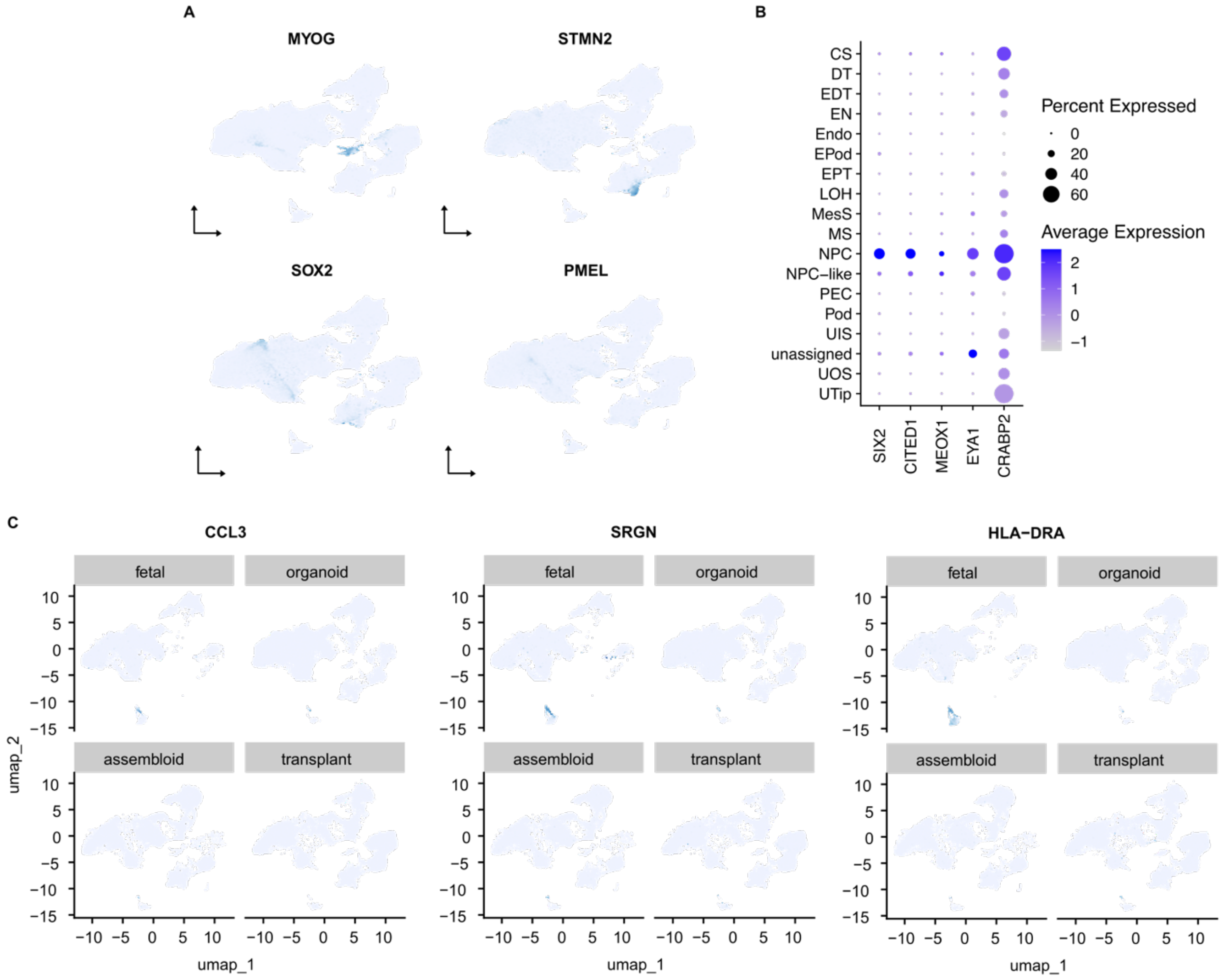


**Figure S1, related to Figure 2. Unassigned, off-target and immune populations in fetal, organoid, assembloid and transplanted kidney tissue. A**) Expression of off-target muscle marker MYOG, neural markers STMN2 and SOX2, and melanocyte marker PMEL. **B**) Expression SIX2, CITED1, MEOX1, EYA1, and CRABP2 across annotated clusters. **C**) Immune markers CCL3, SRGN and HLA-DRA across groups.

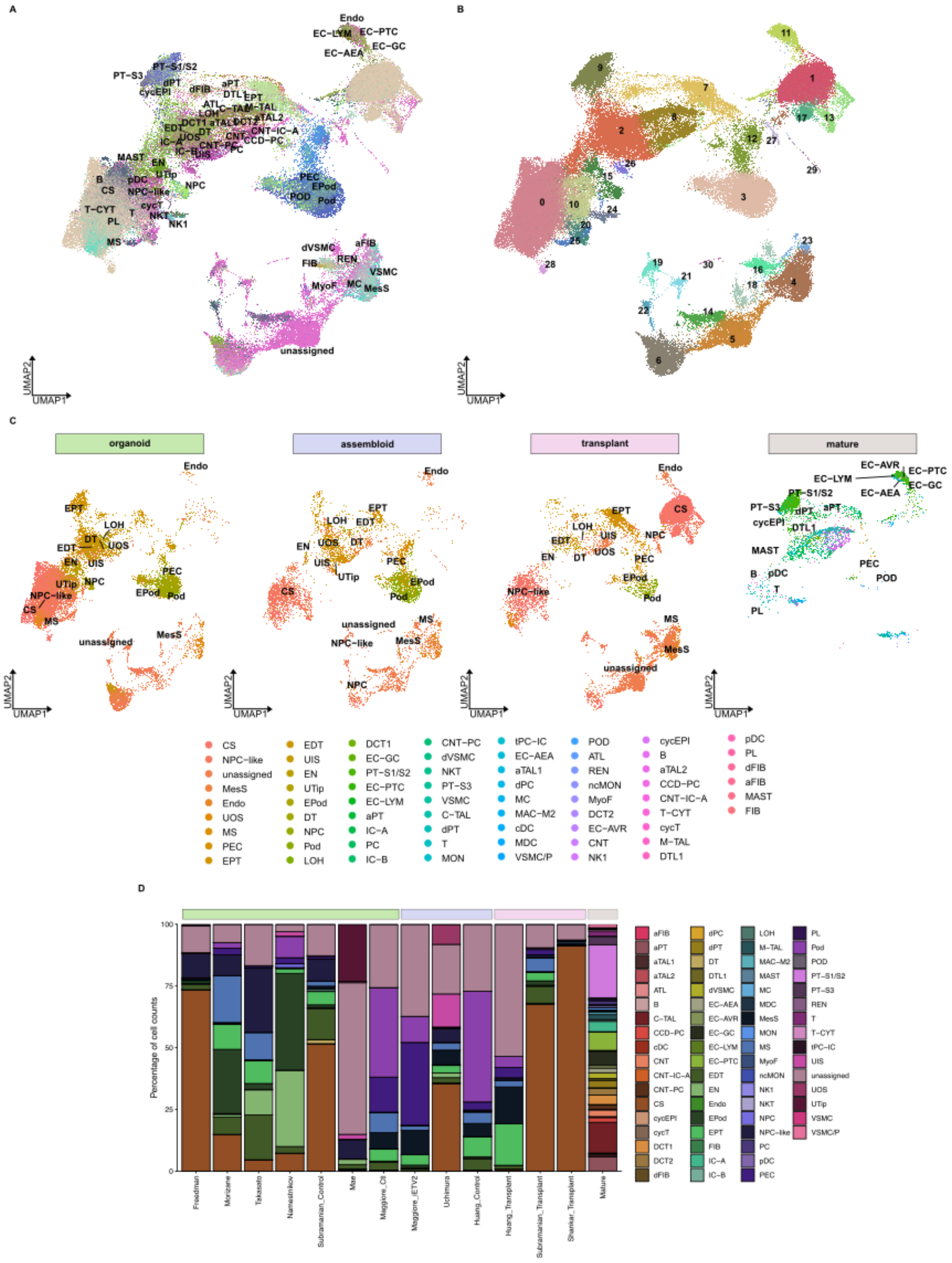


**Figure S2, related to Figure 3. Integration of organoid, assembloid, transplant, and mature kidney tissue.** UMAPs clustered by combined cell type (DKCC+KPMP labels; **A**) and seurat clusters (**B**). **C**) Cell type distribution by group: organoid, assembloid, transplant, and mature tissue. **D**) Cell type proportions by protocol

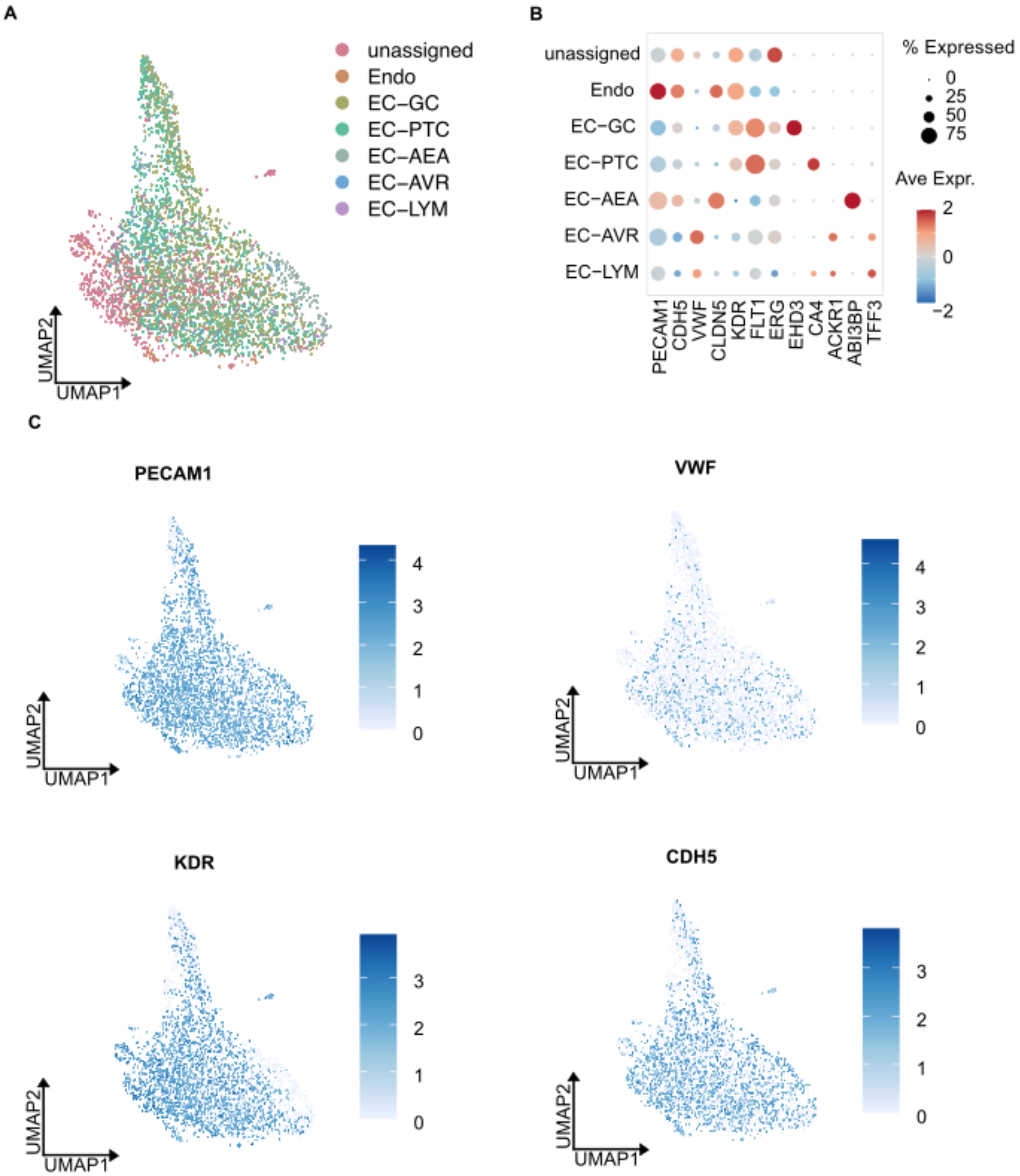


**Figure S3, related to Figure 3. Endothelial population reclassification. A**) Cluster 11 distribution of annotated cell types. Expression of key endothelial markers PECAM1, CDH5, VWF, KDR and FLT1 across mature endothelial subtypes (EC-GC, EC-PTC, EC-AEA, EC-AVR, EC-LYM) and non-mature tissue (Endo and unassigned) (**B**), and within cluster 11 (**C**).

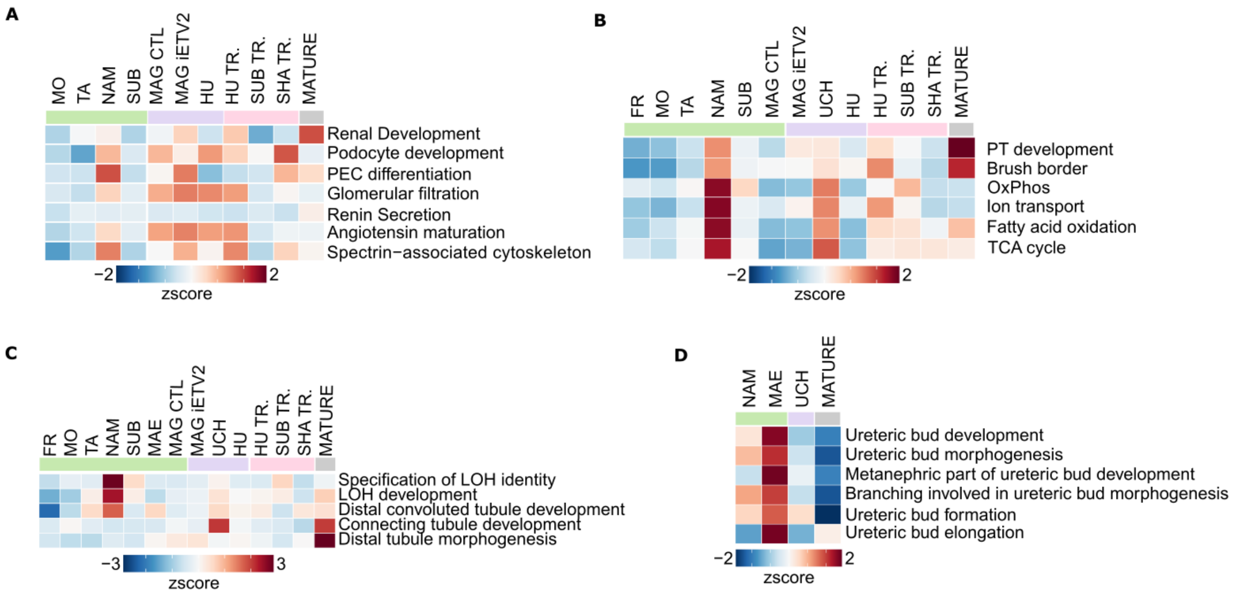


**Figure S4, related to Figure 3. Gene Ontology (GO) enrichment analysis of nephron and ureteric epithelial compartments across protocols.** Heatmaps displaying z-scored AUCell enrichment scores for GO biological process terms associated with renal corpuscle (**A**), proximal nephron (**B**), distal nephron (**C**), and ureteric epithelium (**D**) identity and function. Each row represents a GO term and each column represents a protocol. Enrichment was calculated per cell and aggregated across protocols.

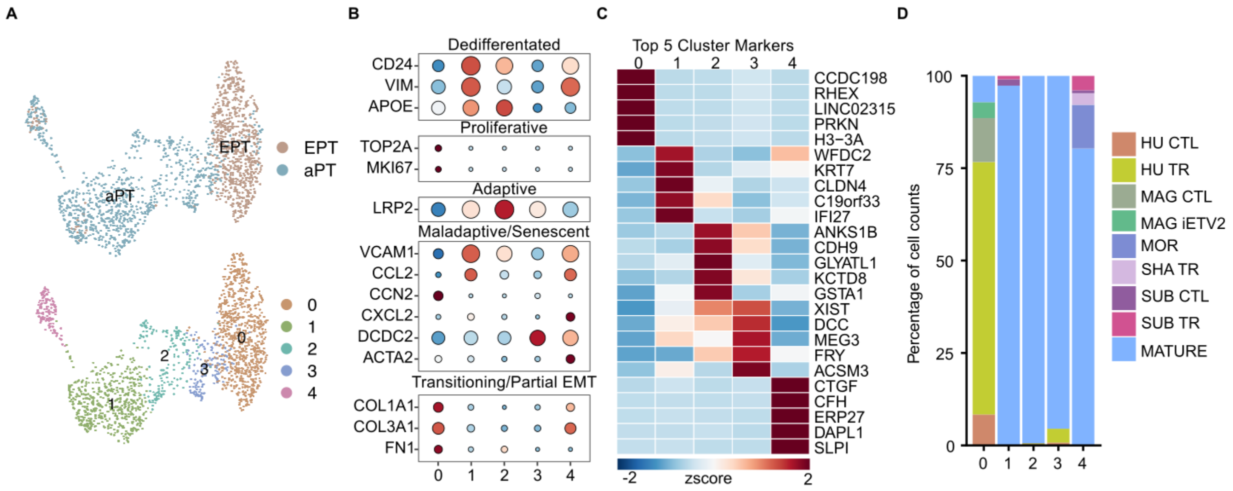


**Figure S5, related to Figure 3. Subclustering of adaptive (aPT) and early proximal tubule (EPT) populations. A**) UMAP of subclustered aPT and EPT populations colored by cluster. **B**) Expression of key marker genes across subclusters, including dedifferentiating markers (CD24, VIM, APOE), maladaptive markers (VCAM1, CCL2, CXCL2, DCDC2), adaptive markers (LRP2), EMT/ECM markers (COL1A1, COL3A1, FN1), and proliferative markers (TOP2A, MKI67). **C**) Top 5 enriched genes across clusters. **D**) Cluster proportion by protocol.

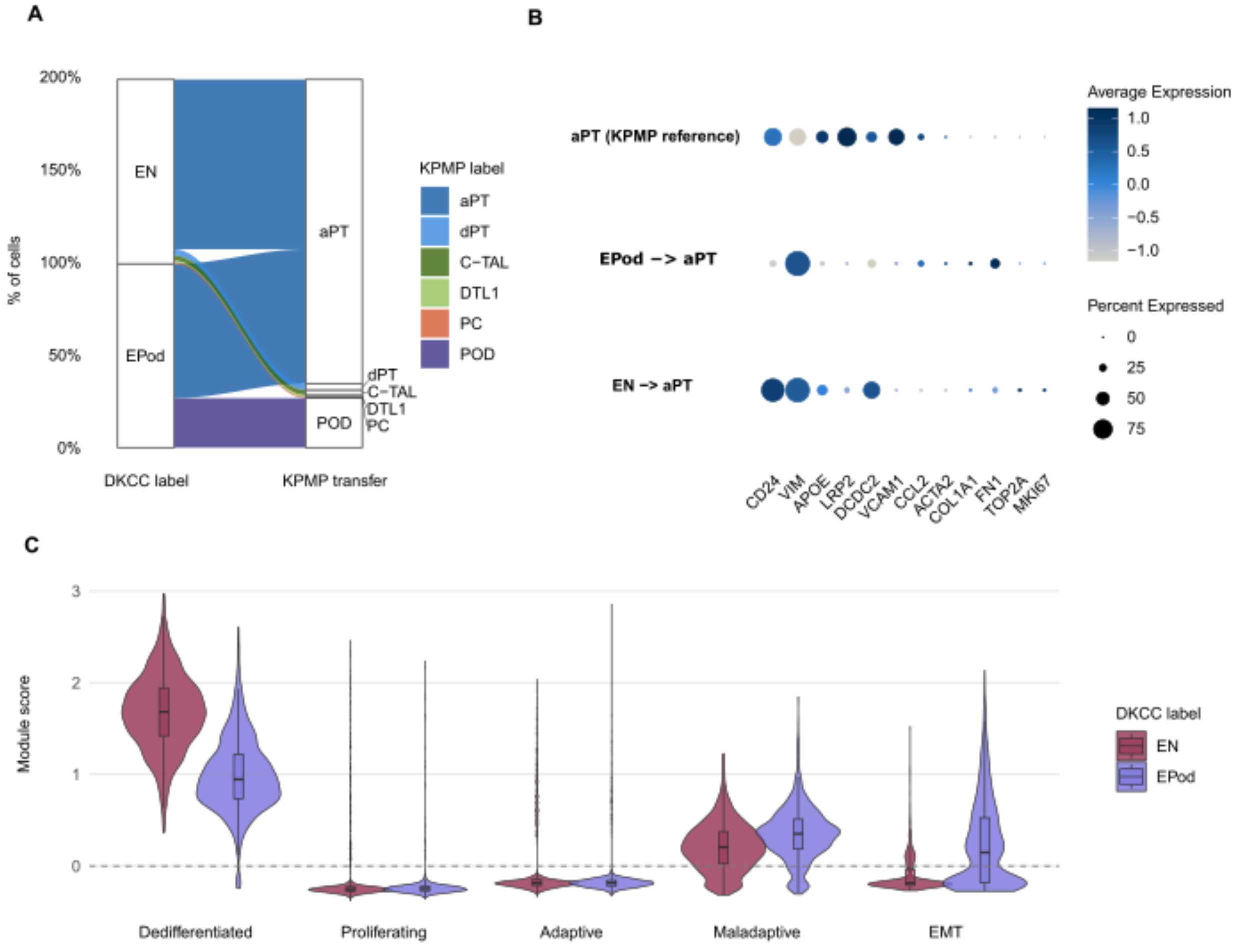


**Figure S6, related to Figure 4. Namestnikov hFKO progenitors map to dedifferentiated adaptive proximal tubules in the KPMP adult kidney atlas. A**) Distribution of KPMP label transfer predictions for EN and EPod in Namestnikov hFKOs. Width is proportional to percentage of cells mapping to each KPMP cell type. **B**) Expression of aPT marker genes across EN and Epod cells mapping to aPT, compared to genuine aPT cells from the KPMP adult kidney reference. **C**) Module scores for dedifferentiated (CD24, VIM, APOE), adaptive (LRP2), maladaptive (DCDC2, VCAM1, CCL2, ACTA2), parital EMT (COL1A1, FN1), and proliferating (TOP2A, MKI67) transcriptional programs in EN and EPod cells mapping to aPT. Boxes show interquartile range.
